# Hypoxic-Core (HyCo) Spheroids Recapitulate Hallmarks of Clinical Hypoxia: A Simple Chip-Based Method for Translational Oncology

**DOI:** 10.1101/2025.04.03.647075

**Authors:** Elena Refet-Mollof, Rodin Chermat, Julie Lafontaine, David (Run Zhou) Ye, Thomas Gervais, Philip Wong

## Abstract

Hypoxia influences the biology and response of cancers. No user-friendly device allows the study of hypoxia on highly-controlled clinically-relevant tumor models. Here, we describe how hypoxic-core (HyCo) spheroids generated using our unique non-perfused microfluidic device recapitulate key clinical hallmarks of hypoxia *in vitro*. Our PDMS-made system can generate up to 240 spheroids naturally exhibiting a diffusion-driven hypoxic core in only 4 days, here from two sarcoma cell lines. Compared to smaller normoxic spheroids from the same cell lines, known hypoxia-related genes are upregulated in HyCo spheroids. In addition, HyCo spheroids display hallmark hypoxia-induced resistance to radiotherapy and chemotherapy, along with increased invasiveness. Finally, to demonstrate applications of our HyCo spheroids and on-chip framework in drug development, we used our HyCo-derived gene expression dataset to select a drug candidate (diethyl-pythiDC) and confirmed its effect on spheroid invasiveness. Our results suggest that HyCo spheroids can be efficiently used as translational tools to integrate hypoxia in cancer research, without complex workflows or setups.

**Teaser:** On-chip spheroids with a natural hypoxic-core emulate clinical cancer hypoxia hallmarks essential for novel therapy development.

## Introduction

Tumor hypoxia refers to intratumoral oxygenation lower than physiological levels and is known to directly increase tumor aggressivity, treatment resistance, metastasis progression and shorten cancer patient survival (1–3). Cellular response to hypoxia involves activation of the hypoxia response pathway, a cascade of physiological and biological changes allowing hypoxic cells to survive in extreme conditions of low oxygen, low pH, and low concentration of nutrients (4–6). These changes include increased glycolysis, angiogenesis, pH regulation, and endothelial to mesenchymal transition (6). Moreover, the hypoxic microenvironment promotes tumor progression and metastasis by exacerbating cell motility and propensity to invade local tissue (i.e., the first step of the metastatic cascade) through both autocrine and paracrine mechanisms (7–9). These mechanisms include secretion of growth factors and modulation of the expression of cell-cell or cell-extracellular matrix adhesion molecules (10). In addition, hypoxia fosters pro-migratory and invasive phenotypes as well as regulating cell intravasation and extravasation, further contributing to metastasis progression (9,10). These combined effects will establish a vicious circle, as the increased tumor volume will result in greater depletion of oxygen and nutrients, further driving the aggressive phenotype (10).

Notably, hypoxia also contributes to anti-cancer treatment resistance (10–13). For example, radiotherapy (RT) efficacy is based on the generation of free radicals within the cell, among which the hydroxyl radical OH•, in turn producing radicals on the DNA (DNA•) (14). As the fixation of these radicals into permanent DNA damage is an oxidative process, hypoxia directly results in decreased RT efficacy (1,14,15). In addition, hypoxic cancer cells have acquired the ability to survive both RT-induced DNA damages and their extensive baseline genomic instability resulting from high cellular stress and impaired capacity for DNA-repair, further limiting the impact of RT (11). Similarly, chemotherapies (CT) that target DNA are less efficient in hypoxic conditions (10,11,16,17). Furthermore, hypoxia increases the expression of a panel of genes involved in CT drug efflux (ATP-binding cassette family, e.g., ABCB1, ABCC1) thereby reducing intracellular drug concentration (18,19). Finally, similar observations can be made regarding immunotherapy, with hypoxia driving an immunosuppressive phenotype (20). Mainly, immune cells cannot easily penetrate hypoxic regions, and the few cells that manage to do so are inactivated due to the acidity of the tumor microenvironment (20,21).

Soft-tissue sarcoma (STS) is a rare groups of cancers in adults (less than 1%) but accounts for 20 % of pediatric solid malignant cancer (22,23). Originating from any soft tissues (e.g., muscle, fat, blood vessels, nerves, tendons, linings of the joints), about 45% of STSs harbor hypoxia, which is associated with worsened clinical outcome (24–26). RT and limb-sparing surgical resection are the standard of care for most large high-grade STS, with RT being preferentially given before surgery (i.e. neoadjuvant RT) to limit long-term RT-related toxicities (27). The value of adjuvant CT is questionable, with no significant differences observed in either relapse-free survival or overall survival (OS) (28). Despite achieving excellent local tumor control through neoadjuvant RT + surgery, around half of patients subsequently develop metastases and their OS is often measured in months (29). For metastatic patients, anthracycline CT is considered standard of care although response rates are extremely poor (e.g., ∼20% for doxorubicin), and is sometimes combined with the alkylating agent ifosfamide (30–32). Cisplatin, another widely-used alkylating agent, has shown poor efficacy in metastatic STSs. However multiple clinical trials using cisplatin in combination with other drugs are still ongoing (NCT06684327, NCT05057130, NCT04605770) (32). As previously stated, the hypoxic nature of STSs likely renders them resistant to CT and radiotherapy, more aggressive and more metastatic, explaining poor patient survival rates (33–35). As such, the validation of alternative treatment strategies taking hypoxia into account is required to reduce metastasis occurrence in STSs patients and increase disease-free survival.

However, there is an apparent lack of user-friendly devices allowing the study of tumor hypoxia in a controlled manner (36–39). Indeed, in vitro study of hypoxia often involves complex experimental setups to control the oxygen level either physically (e.g., hypoxia chambers) or chemically (e.g., cobalt chloride) (36,40). Generally speaking, physical methods do not allow long-term manipulation and RT/drug treatment of samples without removing the samples from the setup, thereby re-oxygenating them. Additionally, oxygen level in such setups is controlled by injecting nitrogen, the resulting media acidification having been shown to induce uncontrolled biases in biology experiments (14,39,41). Similarly, chemical induction can interact with other metabolic pathways than response to hypoxia and eventually affect treatment response evaluation (42). Furthermore, none of these methods are able to capture the whole picture of clinical hypoxia, among which the complex interplay between hypoxic and normoxic cells within the same sample (36,43). Finally, discrepancies between in vitro methods of hypoxia induction limits the establishment of clinically-translatable signatures (44). If in vivo tumor models can naturally exhibit hypoxia, duration of experiments is drastically limited by the need for these tumors to already be extremely large to be sufficiently hypoxic, close to the limit accepted by most ethics committees. Moreover, in vivo quantification of hypoxia is either a post-mortem process or requires complex imaging techniques and equipment (45). As such, despite higher biological relevance, animal models are ill-suited for the study of hypoxia in early preclinical studies.

Microfluidics have eased cancer translational research and drug development for the past 20 years, enabling the culture and treatment of increasingly complex 3D tumor models in compact user-friendly systems (38,46). As the recently signed FDA Modernization Act 2.0, and the proposed FDA Modernization Act 3.0, allow and encourage “cell-based assays, microphysiological systems, or bioprinted or computer models” as alternatives to animal testing for purposes of drug and biological product development, the validation of microfluidic tools is both warranted and timely (47,48). In 2021, we reported a user-friendly non-perfused microfluidic chip on which users can culture up to 240 spheroids that naturally harbor both a normoxic shell and a hypoxic core (49). In this study we delved deeper into the characterization and demonstration of translational applications of our in vitro models: the Hypoxic-Core (HyCo) spheroids. We show that our spheroid model displays key hallmarks of *in vivo* hypoxic STSs, including hypoxia-specific gene expression, treatment resistance and increased invasiveness. Then, we showcase how our on-chip model and methodology can be applied to data-driven treatment testing. Overall, we show that our on-chip tumor model provide cancer researchers with a simple yet clinically-relevant and powerful tool to study hypoxia and its impact on cancer treatment, as well as identifying potential prognostic biomarkers or druggable targets.

## Results

### HyCo spheroids generated on-chip overexpress key known hypoxia-related genes

Our microfluidic chips (Fig. 2A) are transparent, oxygen-permeable, radio-compatible, and allow formation of HyCo spheroids within 2 to 3 days for both SK-LMS-1 and STS117 cell lines. We selected 950 µm as the target diameter of HyCo spheroids based on our in-house in silico model of oxygen consumption (49). At this size, approximately 43% of the spheroid volume or cellular content would theoretically grow under the 10 mmHg pO2 threshold for CAIX-expression, ergo in hypoxia (Fig. 2B, Fig. S1). For SK-LMS-1, HyCo and normoxic spheroids harbor a mean diameter of 921 ± 90 µm and 446 ± 49 µm, respectively (Fig. 2C, D). For STS117 HyCo and normoxic spheroids harbor a mean diameter of 946 ± 75 µm and 465 ± 43 µm, respectively (Fig. 2E, F). It is worth noting that very few HyCo spheroids (3 SK-LMS-1 and 1 STS117, across 38 and 33 spheroids over 3 experiments) are much smaller than average and overlap with normoxic spheroid size distribution. In such cases, “small” HyCo spheroids were excluded from further experiment to prevent confounding effects. As such, as shown in Fig. 2G, our on-chip HyCo spheroids consistently exhibit a CAIX-expressing hypoxic core that occupies approximately 60 % of their cross-section. By extrapolating to the third dimension (Fig. 2B), our results indicate that approximately 45% of the total amount of cells in a HyCo spheroid are hypoxic, consistent with our in silico modeling (Fig. S1).

**Fig. 1.**
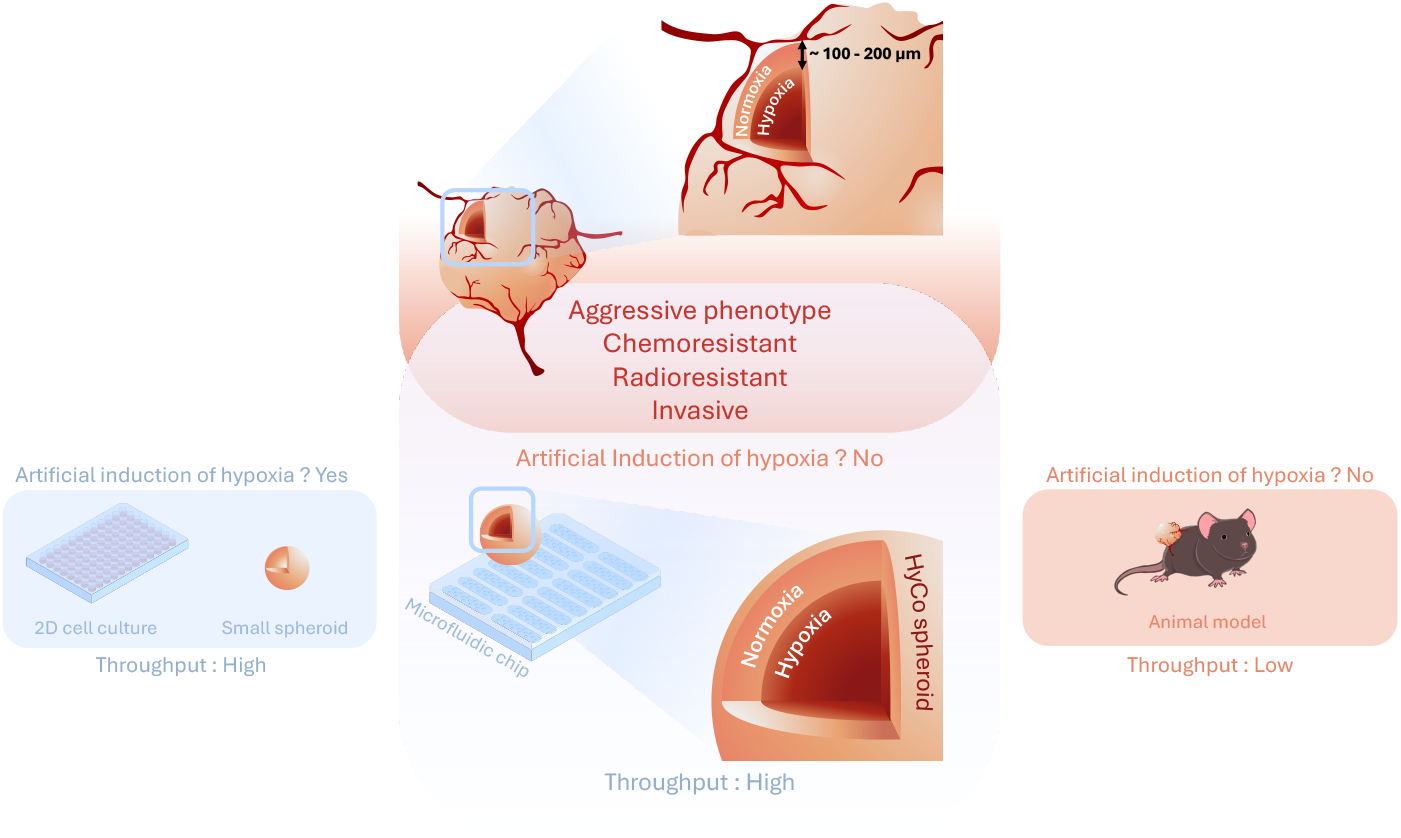
Paradigm for on-chip HyCo spheroids. Soft-tissue sarcomas are hypoxic solid tumors, which contributes to their agressive phenotype, chemoresistance, radioresistance and increased invasiveness. On one hand, in vitro models are high-throughput but require artificial induction. On the other, animal models are naturally hypoxic but are low-throughput and cumbersome. In this study, we showcase how our chip-based high-throughput method generates naturally-hypoxic HyCo spheroids which recapitulate clinical hallmarks of tumor hypoxia, making them adapted for early preclinical studies.

**Fig. 2.**
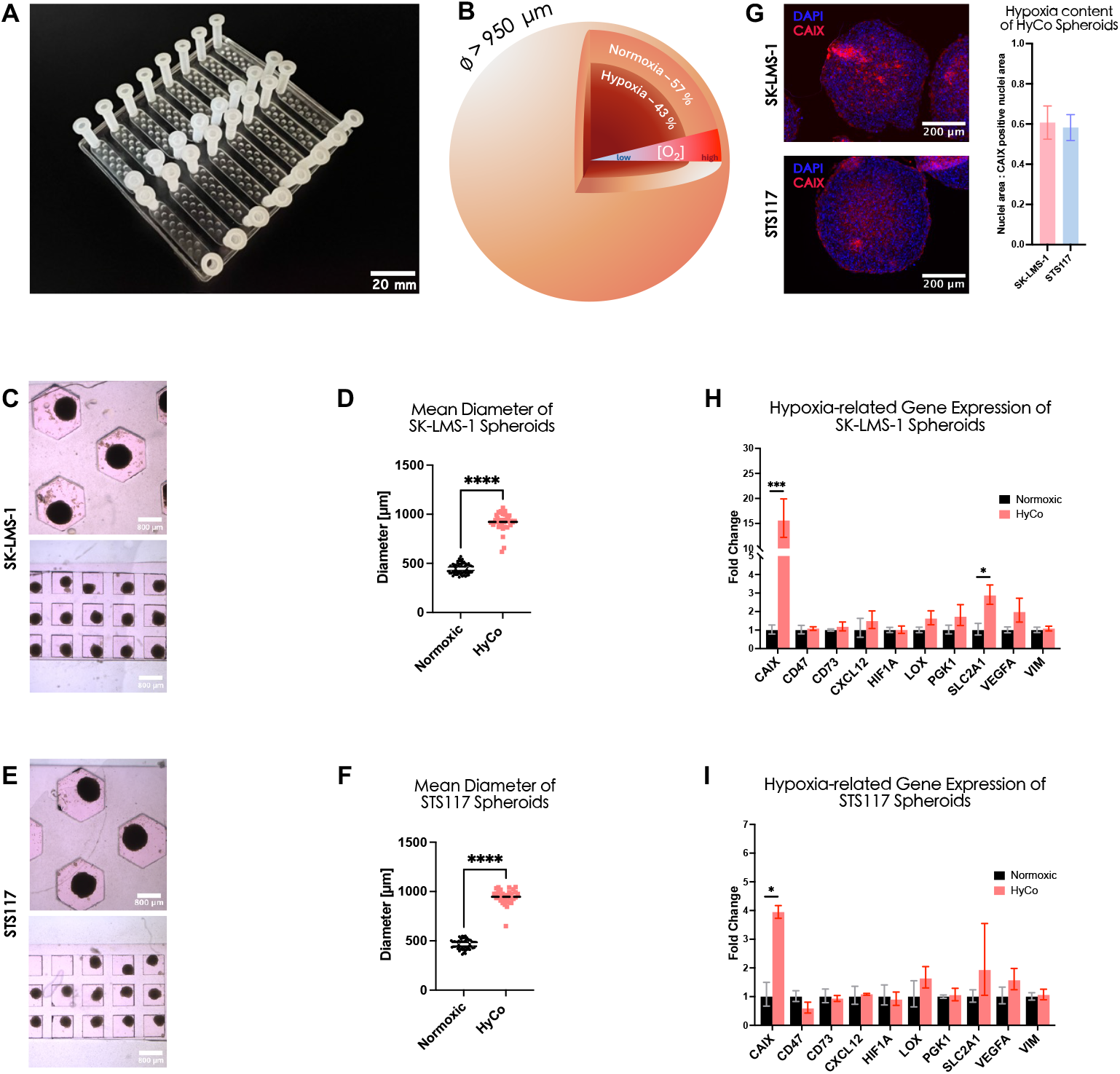
HyCo spheroids have an increased expression of hypoxia-related genes, for both SK-LMS-1 and STS117. (A) Photograph of microfluidic chip for HyCo spheroid generation, culture and treatment. (B) 3D Schematic of HyCo spheroid with sectional view representing the 70:30 normoxic to hypoxic cells volumetric ratio. (C), (E) Brightfield images of SK-LMS-1 and STS117 HyCo and normoxic spheroids. (D), (F) Mean diameter of SK-LMS-1 and STS117 HyCo and normoxic spheroids (**** : p-value < 0.00001, N = 3 independent experiments, total of n= 33 to 79 spheroids, unpaired t-test). (G) Immunofluorescence images of SK-LMS-1 and STS117 HyCo spheroids stained for nuclei (DAPI, in blue) and hypoxia (CAIX, in red), along with quantification of CAIX-positive nuclei area ratio. (H), (I) RTqPCR of hypoxia-related genes expression (CAIX, CD47, CD73, CXCL12, GAPDH, HIF1A, LOX, PDK1, SCL2A1 VEGFA, VIM) in SK-LMS-1 and STS117 HyCo vs normoxic spheroid (mean ± SEM, N = 3 to 4 independent experiments).

To probe wether ∼ 45 % of hypoxic spheroid content was sufficient to induce observable downstream molecular responses, we performed a RTqPCR on a panel of 10 genes known to be upregulated under hypoxic conditions : CAIX, CD47, CD73, CXCL12, HIF1A, LOX, PGK1, SLC2A1, VEGFA and VIM. In both cell lines, most genes of the selected hypoxia-related panel are upregulated (fold-change > 1) except for CD47 in STS117 (Fig. 2H, I). For SK-LMS-1, CAIX (p-value = 0.02), CXCL12 (p-value = 0.21), LOX (p-value = 0.11), PGK1 (p-value = 0.23), SCL2A1 (p-value = 0.03) and VEGFA (p-value = 0.13) have a fold-change above 2 in HyCo spheroids. For STS117, CAIX (p-value = 0.04), LOX (p-value = 0.15), SCL2A1 (p-value = 0.31) and VEGFA (p-value = 0.19) have a fold-change above 2 in HyCo spheroids. Of note, HIF1A gene is not significantly upregulated, with a fold-change of only 1.35 ± 0.62 for SK-LMS-1 and 1.07 ± 0.22 for STS117. Absence of HIF1A upregulation despite evidence of downstream effects of HIF1*α* stabilization has been previously reported for multiple cancers, and is observed here for our two sarcoma cell lines (50).

Overall, HyCo spheroids had increased expression of an array of known hypoxia-related genes in comparison to smaller normoxic spheroids, for both cell lines.

### HyCo spheroids gene expression matches known soft-tissue sarcoma hypoxia signature

Following our RTqPCR, we profiled the bulk transcriptomes of HyCo and normoxic spheroids for SK-LMS-1 and STS117 (Fig. 3). As shown in the first Volcano plot (Fig. 3A), 112 genes are significantly upregulated (log2 fold change > 0.5, p-value < 10e-6), and 43 genes are significantly downregulated (log2 fold change < -0.5, p-value < 10e-6) for SK-LMS-1 HyCo spheroids. As shown in Fig. 3B., only 10 genes are significantly upregulated (log2 fold change > 0.5, p-value < 10e-6), and 3 genes significantly downregulated (log2 fold change < -0.5, p-value < 10e-6) for STS117 HyCo spheroids. Combined analysis of SK-LMS-1 and STS117 data identified shared upregulated genes for future experimental purposes (Fig. S2).

**Fig. 3.**
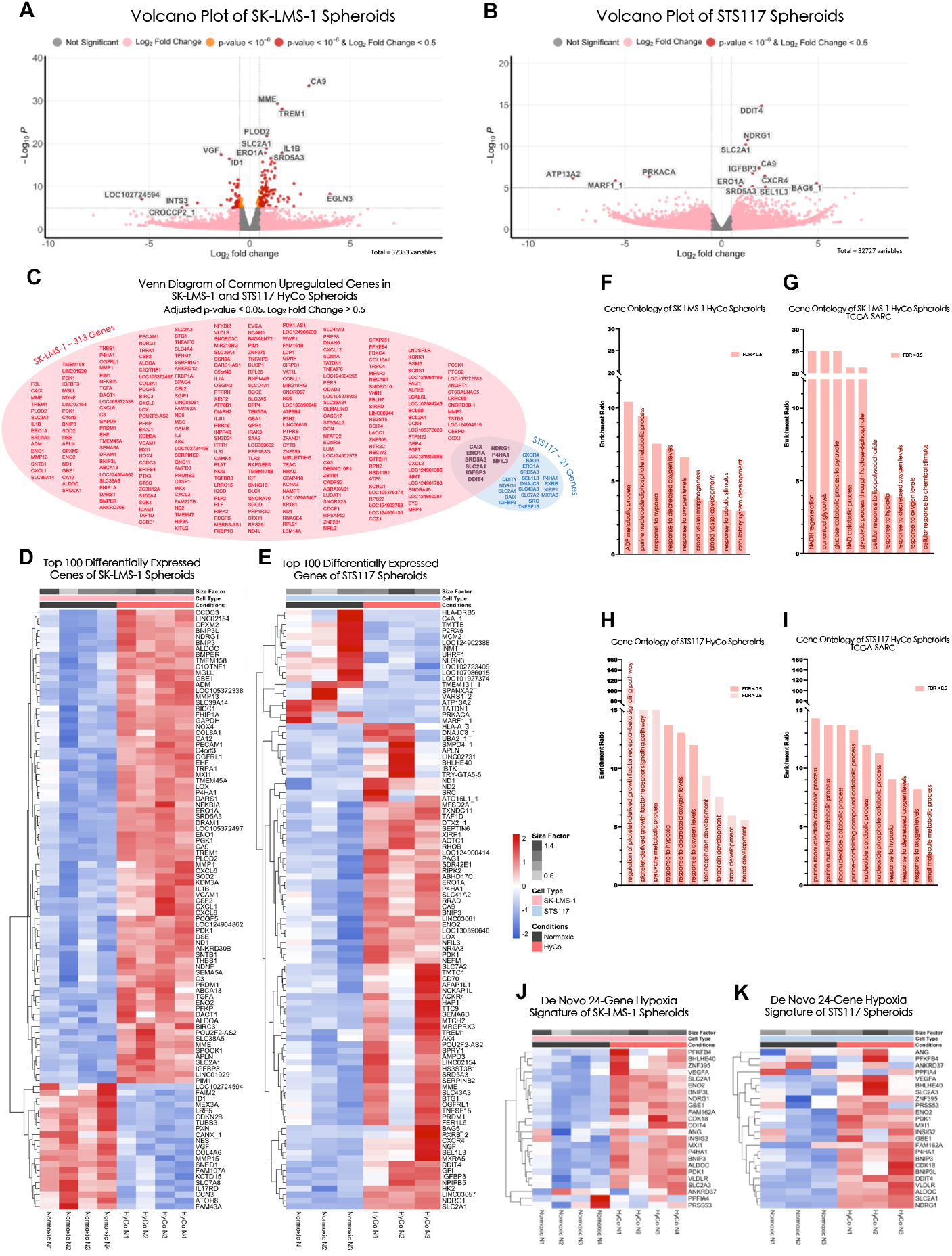
HyCo spheroids hypoxia-related genes signature correlates with STSs clinical data, for both SK-LMS-1 and STS117. (A), (B) Volcano Plot of SK-LMS-1 and STS117 HyCo spheroids compared to normoxic spheroids. (C) Venn diagram of common upregulated genes in both SK-LMS-1 and STS117 HyCo spheroids (adjusted p-value < 0.05, Log2 Fold Change > 0.5). (D), (E) Heatmap of top 100 differentially expressed genes of SK-LMS-1 and STS117 HyCo and normoxic spheroids. (F), (G) Gene ontology pathway enrichment ratio of SK-LMS-1 HyCo spheroids with whole human genome for reference and with TCGA-SARC for reference. (H), (I) Gene ontology pathway enrichment ratio of STS117 HyCo spheroids with whole human genome for reference and with TCGA-SARC for reference. For gene ontology analysis, False Discovery Rate (FDR) correction is applied using Benjamini-Hochberg (BH) method. (J), (K) Heatmap of de novo 24-gene hypoxia signature of SK-LMS-1 and STS117 HyCo and normoxic spheroids. All Heatmaps are clustered by Complete linkage with Euclidean distance and gene expression data are scaled by row (Z-score transformation) before clustering. All data comes from N = 3 to 4 independent experiments.

For each cell line, we then ranked upregulated genes (log2 fold change > 0.5) by their adjusted p-values. As shown in Fig. 3C, out of the 313 and 21 genes that were significanlty upregulated (adjusted p-value < 0.05) in SK-LMS-1 and STS117 respectively, there were 9 genes upregulated in both cell lines: CAIX, ERO1A, SRD5A3, SLC2A1, IGFBP3, DDIT4, NDRG1, P4HA1 and NFIL3.

To visualize gene expression across our samples, we generated heatmaps using the expression levels of the top 100 differentially expressed genes, for SK-LMS-1 and STS117 respectively (Fig. 3D, E). For SK-LMS-1, Unsupervised clustering of the spheroids based on the top 100 differentially expressed genes showed a clear divide between normoxic and HyCo spheroids in both cell lines, with 21 genes downregulated and 79 upregulated in the latter. Similarly, for STS117, 19 genes are downregulated and 81 are upregulated in HyCo spheroids. Based on whole human genome database (RampDB Genomics), gene ontology (GO) terms enriched in both SK-LMS-1 and STS117 HyCo spheroids include pathways in *response to hypoxia, response to oxygen level* and *response to decrease oxygen levels*. (Fig. 3F, H). Other GO terms enriched in SK-LMS-1 are related to metabolism switch and vasculogenesis, of which the relationship with hypoxia has been well-documented (9,36). For other pathways enriched in STS117 however, FDR are consistently above significance threshold and pathways such as *brain development* and *forebrain development* are, at first glance, unrelated to STSs response to hypoxia. Although the biological relevance of this finding cannot be entirely ruled out, considering the genome as a whole could lead to probability-based biases and misinterpretation of enriched pathways. Therefore, we performed gene ontology analysis using the human sarcoma-specific database (TCGA-SARC) as a reference to help us interpret our findings in the context of sarcomas. Based on TCGA-SARC database, GO terms enriched in HyCo spheroids of both cell lines now only include pathways linked to low oxygen levels and metabolic adaptation to hypoxia, in accordance with literature (Fig. 3G, I) (51).

Finally, we applied the *de novo* 24-gene hypoxia signature from Yang et al. to our data, with expression levels for SK-LMS-1 and STS117 spheroids illustrated in Fig. 3J and 3K (51). Out of the 24 genes, 15 and 18 are upregulated (log2 fold change > 0.5) within HyCo spheroids for SK-LMS-1 and STS117 respectively (Fig. 3J, K). This showcases that HyCo spheroids gene signature matches that of Yang et al., which was selected based on its specificity to hypoxic STSs and its correlation with shorter pastient distant metastasis-free survival (51).

Overall, these results confirm the hypoxic nature of our HyCo spheroid model and its biological relevance in preclinical research.

### HyCo spheroids recapitulate clinically-observed hypoxia-induced treatment resistance

To further confirm the hallmark phenotypical behavior and more precisely the resistance to both chemo- and radiotherapy of our HyCo spheroids, we treated them with 5 µM of cisplatin or with 4 and 8 Gy external beam radiotherapy (EBRT). Cisplatin, a known alkylating chemotherapeutic agent, induces DNA-damages ultimately leading to apoptosis. Therefore, we evaluated cytotoxicity by quantifying accumulation of caspase 3 and 7 activation over 16 h. We selected cisplatin specifically for its poor response rate in STSs (10 %) and the well-established links between hypoxia and cisplatin resistance (17,32). Although EBRT also inflicts DNA-damages, we evaluated treatment efficacy by quantifying clonogenic survival to capture the long-term consequences of ionizing radiation on tumor cells.

For both cell lines, cisplatin-treated and non-treated HyCo spheroids display similar level of activated caspase 3 and 7, meaning there is no significant effect of cisplatin (Fig. 4A – D). By contrast, normoxic spheroids have 2.47 ± 0.74 times higher activated caspase 3 and 7 than HyCo (adjusted p-value < 0.05) for SK-LMS-1 (Fig. 4C), and 1.69 ± 0.20 (adjusted p-value < 0.05) for STS117 (Fig. 4D), showcasing that normoxic spheroids are more sensitive to cisplatin than HyCo spheroids of the same cell line. In addition, when separating the response to cisplatin between normoxic and hypoxic regions of HyCo spheroids, we observed that both regions exhibited lower activated caspase 3 and 7 than corresponding regions in normoxic spheroids (Fig. S1). Although not significant, this finding points towards paracrine signalling of cisplatin resistance from the hypoxic to the normoxic regions of HyCo spheroids.

**Fig. 4.**
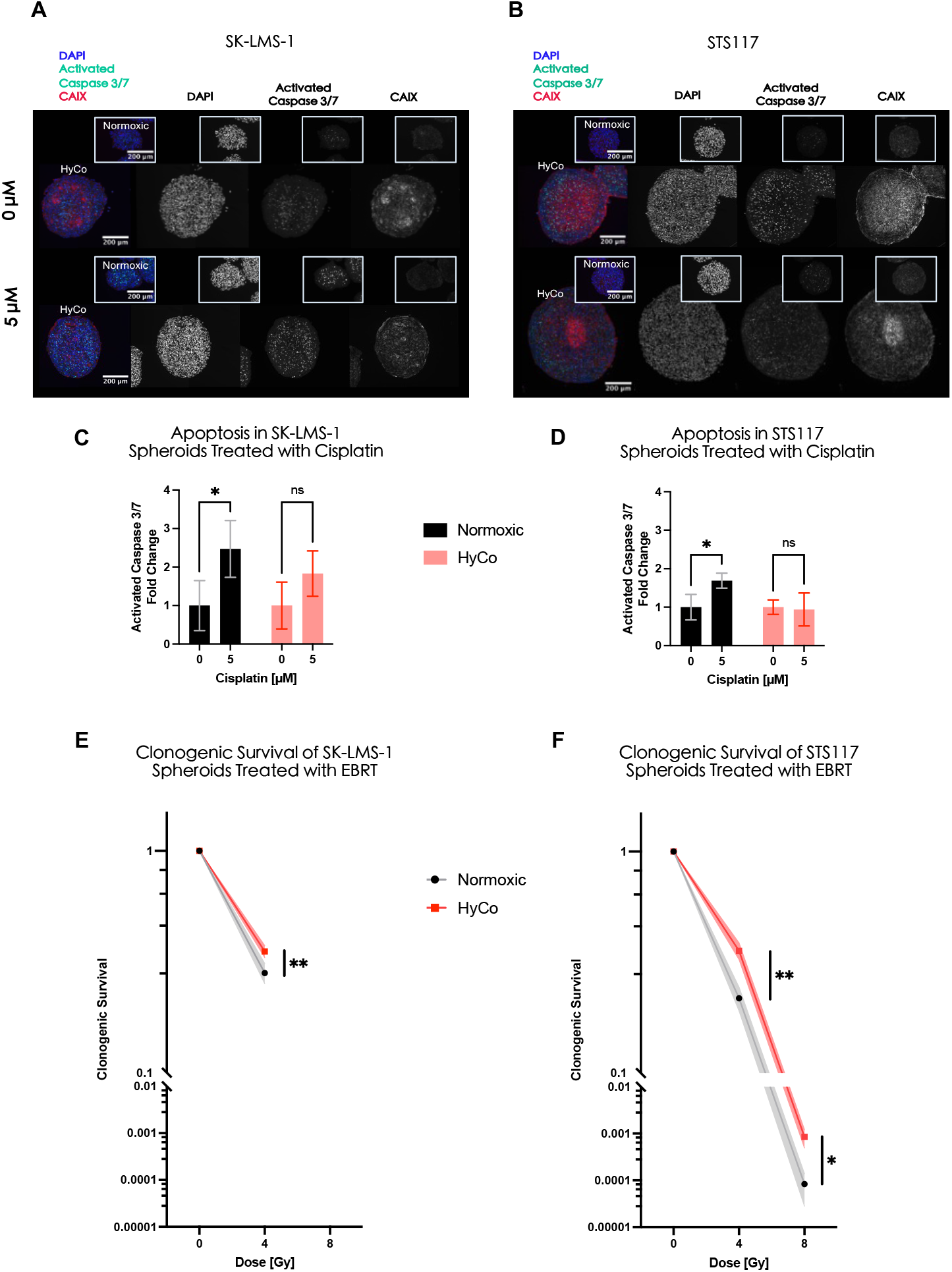
HyCo spheroids are resistant to both chemotherapy and radiotherapy compared to normoxic spheroids. (A), (B) Immunofluorescence images of SK-LMS-1 and STS117 HyCo and normoxic spheroids after overnight exposure to 5µM of cisplatin, stained for nuclei (DAPI in blue), apoptosis (CellEvent ready probe 3/7 in green) and hypoxia (CAIX in red). (C), (D) Quantification of apoptosis signal (activated caspase 3/7) after overnight exposure to 5µM of cisplatin in SK-LMS-1 and STS117 HyCo and normoxic spheroids. Data are presented as mean ± SD, (*) : adjusted p-value < 0.05, N=3,4 independent experiments, n = 2 to 8 spheroids per repetitions, 2way ANOVA, Šidák’s multiple comparisons test . (E), (F) Clonogenic survival curves of SK-LMS-1 and STS117 HyCo and normoxic spheroids after on-chip external beam radiotherapy treatment. Data are presented as mean ± SD, (*) : adjusted p-value < 0.05, (**) : adjusted p-value < 0.01, N=3 independent experiments, multiple unpaired t-test using Holm-Šidák method.

Similarly, the clonogenic assay confirms the resistant behavior of HyCo spheroids for both cell lines after treatment with EBRT. SK-LMS-1 HyCo spheroids exhibit 1.26 ± 0.07 times more colonies than normoxic spheroids after exposure to 4Gy of EBRT (adjusted p-value < 0.01) (Fig. 4E). However, we observed no colonies when treated with 8 Gy of EBRT (Fig. 4E). STS117 HyCo spheroids exhibit 1.64 ± 0.07 times more colonies than normoxic spheroids after exposure to 4 Gy (adjusted p-value < 0.01) and 13.67 ± 8.04 times more colonies than normoxic spheroids after exposure to 8 Gy (adjusted p-value < 0.05) (Fig. 4F). From this data, STS117 cells appear more radioresistant than SK-LMS-1, as they are still able to form colonies after 8 Gy of EBRT even under normoxic conditions. At 4 Gy however, normoxic STS117 spheroids are more sensitive than normoxic SK-LMS-1, while HyCo STS117 and HyCo SK-LMS-1 exhibit similar clonogenic survival. As such, hypoxia-induced radioresistance also appears more prominent for the STS117 cell line at 4 Gy. More broadly, the clonogenic assay indicates a dose response of HyCo and normoxic spheroids to EBRT regardless of cell line.

Overall, for both cell lines, HyCo spheroids are resistant to CT (cisplatin) and EBRT, therefore exhibiting an overall more resistant phenotype compared to their smaller normoxic counterpart.

### HyCo spheroids recapitulate clinically-observed hypoxia-induced tumor agressiveness

Another critical characteristic of hypoxia in cancer is the promotion of an aggressive phenotype, manifesting as increased invasiveness and metastatic progression. To investigate this, we performed an invasion assay on HyCo and normoxic spheroids by following the invasion of spheroids in a Matrigel matrix over 10 days (Fig. 5A). We then analyzed the invasion images using the method described in Fig. 5B, adapted from G. J. Lim et al. (52). Briefly, invasion index A describes the invading outermost regions of the spheroid, while invasion index B includes the loosely aggregating intermediate region (Fig. 5B)

**Fig. 5.**
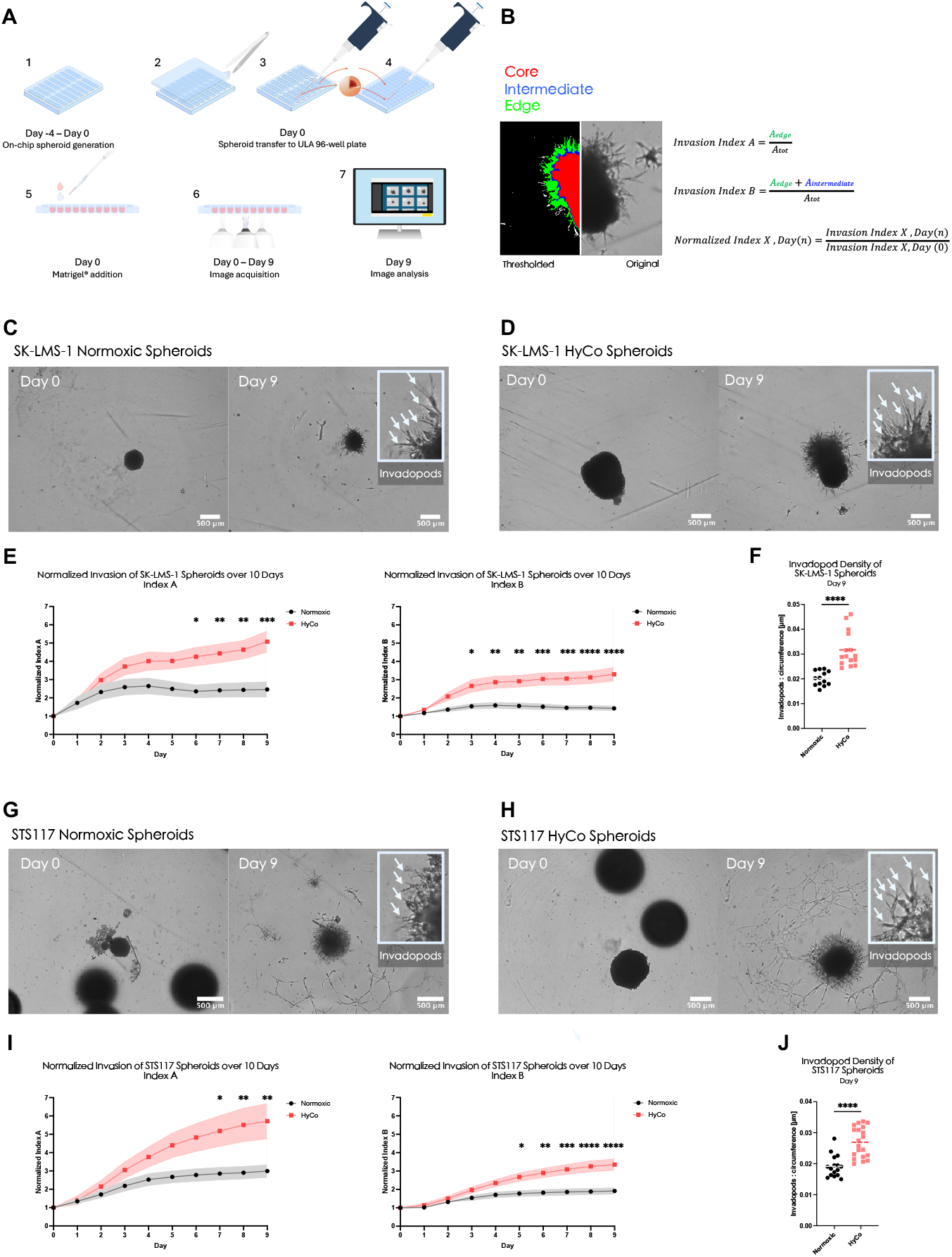
HyCo spheroids are more invasive than small normoxic ones. (A) Schematic of experimental timeline. (B) Example of invasion image analysis pipeline. (C), (D) Brightfield images of SK-LMS-1 normoxic and HyCo spheroids at day 0 and day 9. White arrows indicate invadopods, i.e. cell protrusions into the Matrigel. (E) Normalized invasion indexes A and B for SK-LMS-1 normoxic and HyCo spheroids over 10 days. (F) Invadopod density of SK-LMS-1 normoxic and HyCo spheroids at day 9. (G), (H) Brightfield images of STS117 normoxic and HyCo spheroids at day 0 and day 9. White arrows indicate invadopods, i.e. cell protrusions into the Matrigel. (I) Normalized invasion indexes A and B for STS117 normoxic and HyCo spheroids over 10 days. (J) Invadopod density of SK-LMS-1 normoxic and HyCo spheroids at day 9. All invasion indexes data are presented as mean ± SEM, (*) : adjusted p-value < 0.05, (**) : adjusted p-value < 0.01, (***) : adjusted p-value < 0.001, (****) : adjusted p-value < 0.0001, N = 4 independent experiments, n = 5 normoxic spheroids and n= 5 HyCo spheroids per experiments, 2way ANOVA, Šidák’s multiple comparisons test. Invadopod density data are presented as a scatter plot with mean value, p-value < 0.001, (****), N = 4 independent experiments, n = 5 normoxic spheroids and n= 5 HyCo spheroids per experiments, unpaired t-test.

For both cell lines, normalized invasion indexes A and B were significantly higher for HyCo spheroids compared to normoxic ones (Fig. 5C, D, F). Normalized invasion index A is 1.8-times higher for HyCo spheroids than for normoxic ones on day 7 for SK-LMS-1 (Fig. 5F) and day 6 for STS117 (Fig. 5I) (adjusted p-value < 0.05). On the other hand, normalized invasion index B is 1.7-times higher on day 5 for SK-LMS-1 (Fig. 5F) (adjusted p-value < 0.05) and 1.5-times on day 3 for STS117 (Fig. 5I) (adjusted p-value < 0.05) HyCo spheroids compared to normoxic ones. Furthermore, both normalized indexes remain significantly higher for HyCo spheroids for the remainder of the experiment. It is also worth noting that normalized invasion indexes of HyCo spheroids of both cell lines continuously increase overtime while normalized invasion indexes of normoxic spheroids plateaued starting day 4.

In addition, invasion assays can provide information on the distribution of the various arms protruding from the spheroids into the surrounding Matrigel, which we have termed “invadopods”. As shown on Fig. 5C, D, I, and J, each spheroid is surrounded by dozens of such protrusions, the amount of which can be normalized to the circumference of the invading area to derive an invadopod density. Here, for both SK-LMS-1 and STS117, invadopod density on day 9 is on average higher in HyCo spheroids by 57 % and 40 % respectively (p-value < 0.0001) (Fig. 5G, L). In other words, the proportionally higher invasiveness of HyCo spheroids is also accompanied by a proportionally higher amount of invadopods.

As a side note, across all repetitions, 6 SK-LMS-1 HyCo spheroids underwent a process we labeled “sprouting”, whereby a secondary spheroid is partially or completely expulsed from the primary one (Fig. S4). This process was only observed for HyCo spheroids of the SK-LMS-1 cell line and was not investigated further.

Overall, our results demonstrate that HyCo spheroids exhibit a more invasive phenotype, in accordance with clinical literature on tumor hypoxia.

### HyCo spheroids enable data-driven cancer-specific preclinical drug testing

Finally, we sought to demonstrate applications of our HyCo spheroids to drug testing. Based on genes upregulated in both SK-LMS-1 and STS117 (Fig. 3C), availability of such compounds and documented evidence of preclinical efficacy, we selected P4HA1-specific inhibitor diethyl-pythiDC as a drug candidate. Diethyl-pythiDC mode of action relies on directly decreasing tumor ability to infiltrate local microenvironment, and as such has been shown to reduce tumor invasion and metastasis (53). Given the documented impact of this compound on proliferation and invasion, and given the metastatic nature of STSs, we chose to quantify diethyl-pythiDC efficacy by measuring spheroid invasiveness through invasion assays (Fig. 6A).

**Fig. 6.**
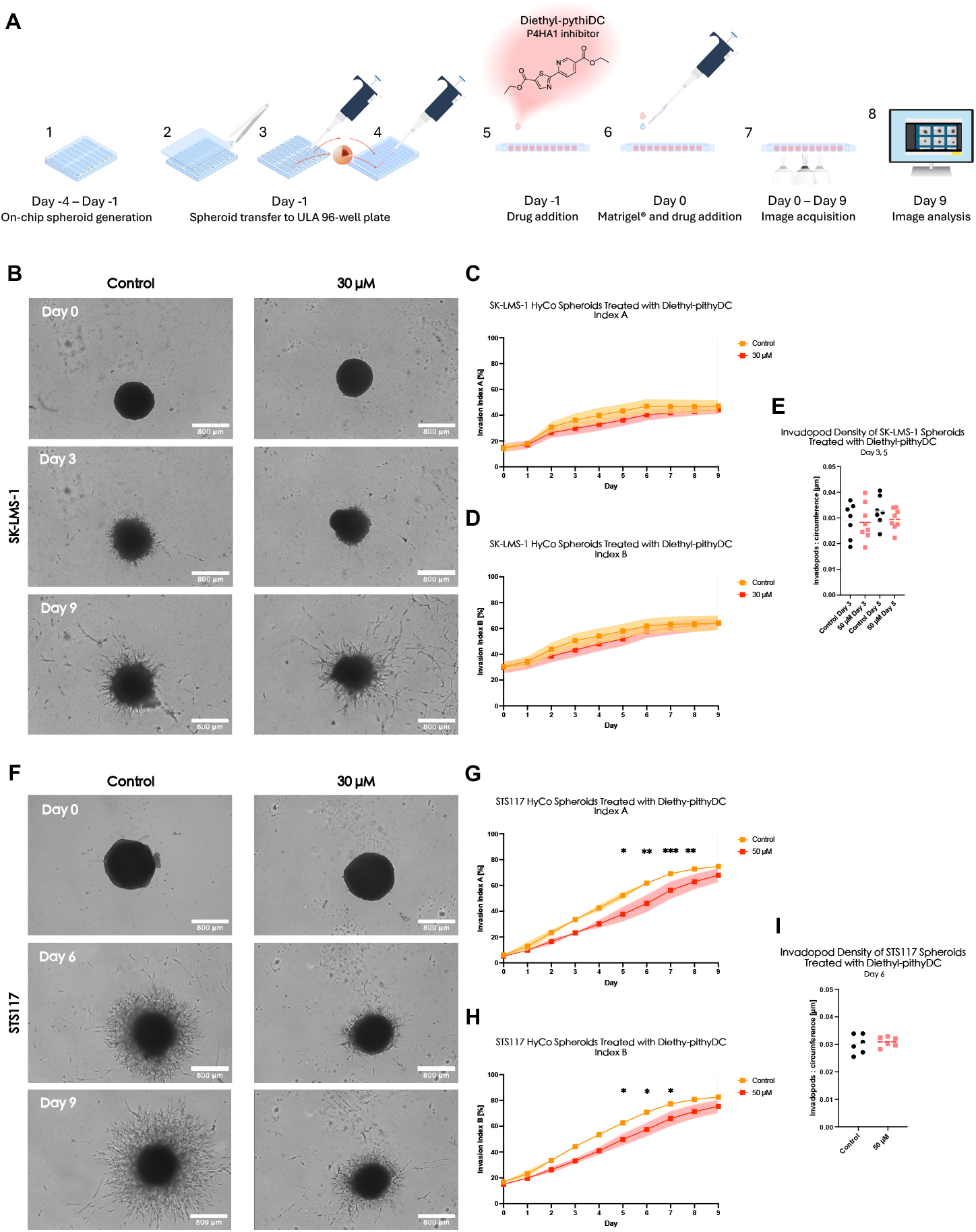
HyCo spheroids can provide preclinical information on drug efficacy. (A) Schematic of experimental timeline. (B) Brightfield images of SK-LMS-1 HyCo spheroids treated with control buffer (left) and diethyl-pythiDC (right) at days 0, 3 and 9. (C),(D) Normalized invasion indexes A and B for SK-LMS-1 control and treated HyCo spheroids over 10 days. (E) Invadopod density of SK-LMS-1 control and treated HyCo spheroids at day 3 and day 5. (F) Brightfield images of STS117 HyCo spheroids treated with control buffer (left) and diethyl-pythiDC (right) at day 0, 6 and 9. (G), (H) Normalized invasion indexes A and B for STS117 control and treated HyCo spheroids over 10 days. (I) Invadopod density of STS117 control and treated HyCo spheroids at day 6. All invasion indexes data are presented as mean ± SEM, (*) : adjusted p-value < 0.05, (**) : adjusted p-value < 0.01, (***) : adjusted p-value < 0.001. N = 4 independent experiments for SK-LMS-1 and N = 3 for STS117. n = 4 HyCo spheroids per experiments (2 control and 2 treated). For SK-LMS-1, a mixed-effects analysis with Šidák’s multiple comparisons test was used due to uneven group sizes (1 blurry spheroid in N3 control group). For STS117, 2way ANOVA, Šidák’s multiple comparisons test analysis was used.

We observed no significant effect of diethyl-pythiDC on SK-LMS-1 HyCo spheroids. Still, both indexes were lower compared to control, with the maximum difference between the 2 conditions observed at day 5 and 3 for indexes A (16.3 %) and B (14.7 %) respectively (Fig 6B-D). Furthermore, we report no differences in invadopod density for spheroids at either day 3 or day 5 (Fig. 6E).

On the other hand, diethyl-pythiDC appears effective for STS117 HyCo spheroids, with indexes A and B being significantly lower for HyCo at days 5 to 8 and 5 to 7 respectively (Fig. 6F-H). This difference is the highest at day 6, with indexes A and B being reduced by 25.4 % (61.8 ± 4.1 to 46.1 ± 16.0, adjusted p-value < 0.001) and 18.9 % (70.9 ± 3.1 to 57.5 ± 12.8, adjusted p-value < 0.05) respectively. However, indexes of the control group appear to ultimately catch up to that of the treated group, with no significant difference at day 9 for index A and at day 8 for index B. Finally, there was no difference in invadopod density between treated and untreated spheroids at day 6 (Fig. 6I). This indicates that the reduced invasiveness is not mediated by reduced number of invadopods but rather by reduced ability of these invadopods to invade, consistent with diethyl-pythiDC mechanism of action.

Overall, these results demonstrate that HyCo spheroids can be efficiently applied to preclinical treatment testing, with their clinically-relevant hypoxia-specific gene signatures being used as guidance for treatment selection.

## Discussion

While tumor hypoxia represents a well-known feature associated with worsened patient survival, there is still no user-friendly platform to study it in laboratory settings. As previously mentioned, hypoxia research requires complex experimental setups such as hypoxic chambers or chemicals (e.g., cobalt chloride), which can introduce biological biases including media acidification and interaction with treatments, assays and outcomes. Conversely, although closer to clinical mechanisms of hypoxia, mice tumor models are heterogeneous in their hypoxic content, both spatially and temporally, thereby increasing the complexity and cost of studying them. Quantification of hypoxia *in vivo* is either post-mortem using immunofluorescence (IF) or immunohistochemistry (IHC) or require methods such as needle-type O2 electrodes or complex and costly imaging techniques (e.g, PET, fMRI), which are not universally available and demand specialized personnel. Overall, animal models, though biologically relevant, are ill-suited to study hypoxia for the first steps of preclinical studies. In 2022, the FDA Modernization Act 2.0 encouraged the use of “cell-based assays, microphysiological systems, or bioprinted or computer models” as alternatives to animal testing for purposes of drug and biological product development (48). A proposal to mandate the FDA to update its regulations within 12 months, dubbed FDA Modernization Act 3.0, has been introduced in the US Senate in January of this year (47). Thus, developing a simple powerful preclinical tools to study hypoxia can contribute to the democratization of scientific research by lowering barriers to entry, encouraging interdisciplinary collaboration, and enabling broader participation in scientific research and hypoxia-related treatment development.

Our microfluidic chips easily produce 240 size-controlled spheroids that naturally harbor both a hypoxic core and a normoxic shell due to oxygen-diffusion limitations, defined as HyCo spheroids. As shown in our previous publication and in Fig.2B and 2C HyCo spheroids reliably feature a hypoxic core comprising 35 - 45% of their total volume (49). In this study, we further characterize > 900 µm spheroids and demonstrated that they upregulate key hypoxia-response genes such as CAIX, SCL2A1, or VEGFA (44). As previously stated, HIF1A gene was not upregulated at the gene level despite demonstrable downstream effects of HIF1*α* protein stabilization, including CAIX expression. Given that most of the volume of HyCo spheroids is in normoxic conditions, it is possible that the signal from hypoxic cells is simply masked by that of normoxic ones (49). Furthermore, although post-translational regulation of HIF1*α* activity under hypoxic conditions is well-known, oxygen-independent regulatory mechanisms of HIF1A transcription and translation have also been identified and observed firsthand in our previous publication (49,50,54).

Although HyCo spheroids are not entirely hypoxic, their gene expressions recapitulate those observed in clinical STSs hypoxic samples. Transcriptomic profiling showed a clear difference in gene expression between HyCo and normoxic spheroids, which was more evident for SK-LMS-1 than STS117. Indeed, SK-LMS-1 HyCo spheroids have 10-times more significantly upregulated genes than STS117 despite having the same controlled hypoxia content. These differences may stem from STS subtype-specific adaptation to hypoxia or inherent differences between commercially-available and patient-derived cell lines, both of which have been documented **(55– 57)**. Genes significantly upregulated in both SK-LMS-1 and STS117 included SLC2A1, SRD5A3, IGFBP3, NDRG1, CAIX and ERO1A, all of which are known to be involved in tumor response to hypoxia at various levels and have been shown to correlate with poorer clinical outcomes (14,42,51).

Our transcriptomic data and GO analyses support the biological relevance and alignment with known clinically-identified hypoxia-related cancer gene expression profiles, and more specifically with STSs-derived profiles (51,58). This solidifies HyCo spheroids as an excellent clinically-relevant in vitro model of STSs hypoxia. Given the heterogeneity of STSs and clinical outcomes, better risk stratification strategies are warranted. Currently, despite the identifications of several prognostic gene signatures, their clinical utility are limited by the lack of efficacious systemic or hypoxia-targeted treatments that could alter patient outcomes (51,58). As HyCo spheroids can be generated from both commercially available and patient-derived cell lines, our chip-based strategy offers a versatile platform for identifying cancer-specific hypoxia biomarker and testing treatment response.

To further validate the functional relevance of our HyCo spheroids, we treated the spheroid on-chip with both CT and EBRT to demonstrate hypoxia related chemo- and radioresistance. We selected cisplatin as it is a widely-used alkylating chemotherapeutic agent, with a 10% clinical response rate in STSs compared to the 20% response rate of doxorubicin, the first-line treatment for STSs (32). Like RT, cisplatin cytotoxicity relies on DNA-damage induction which is an oxygen-dependent process, lessened under hypoxic conditions. In addition, evidence of a variety of hypoxia-related mechanisms of cisplatin resistance (e.g., increased drug efflux, cisplatin detoxification, etc.) have been reported (59). HyCo spheroids recapitulate this phenomenon, exhibiting less apoptosis than their normoxic counter parts when treated with cisplatin. Our results indicate that both normoxic and hypoxic regions of HyCo spheroids display resistance to cisplatin, although the difference between each region and normoxic spheroids is not statistically significant (Fig. S3). This suggests not only a hypoxia-driven chemoresistance in hypoxic cells but also a hypoxia-driven paracrine mechanism of resistance in normoxic cells, both of which have been documented (7,59). Of note SK-LMS-1 were overall more sensitive to cisplatin than STS117. Increased sensitivity to cisplatin can be attributed to a variety of pathways, including the significant downregulation of MYC in SK-LMS-1 HyCo spheroids (60). Additionally, it has been reported that commercial cell lines do not adequately reproduce drug sensitivity in cancer patients, highlighting the value of including patient-derived models in non-clinical drug testing (55,57).

Hypoxia-induced resistance to RT has been widely documented, both in vitro and clinically, and is based on reduced oxygen-dependent fixation of DNA damage (14). The clonogenic assay is the gold-standard for RT efficacy, based on the ability of individual cells to successfully undergo multiple rounds of mitosis after irradiation (61). Here, HyCo spheroids recapitulate the expected radioresistance conferred by hypoxia, as HyCo spheroids have consistently higher clonogenic survival compared to normoxic spheroids. In addition, differential response between SK-LMS-1 and STS117 in both normoxic and HyCo conditions points toward treatment resistance being a function of both oxygen status of the cell as well as sarcoma subtype-dependent factors. As both normoxic and hypoxic cells co-exist within our samples in a controlled manner, performing the clonogenic assay on the whole spheroid can be seen as analogous to evaluating the response of a whole tumor. Once again however, with more than half of the volume of HyCo spheroids in normoxia, the contribution of hypoxic cells to clonogenic survival is significantly diluted. Importantly, unlike hypoxia chambers, our system allows straightforward on-chip irradiation due to its water equivalence and radiocompatibility, enhancing its utility for translational radiotherapy research (46,49,62,63). This further illustrates the usefulness of chip-based approaches for *in vitro* CT and RT testing, especially in the context of translational cancer research.

The invasion assays provided further support for the aggressive hypoxia phenotypes exhibited by HyCo spheroids with significantly increased and continuously increasing normalized invasion indexes compared to normoxic smaller spheroids. Of interest, sprouting event was only observed in HyCo SK-LMS-1 spheroids and warrant future investigation. Contrary to other assays discussed in this manuscript, invasion assays offer a non-destructive way of evaluating whole spheroids, retaining the three-dimensional architecture and the complex interplay between normoxic and hypoxic cells. Yet, the recency of this methodology makes it prone to potential biases in analysis, and metrics should be carefully selected to mitigate them.

In turn, HyCo spheroids are as such uniquely adapted to perform invasion assays, especially in the context of pre-clinical drug development. To demonstrate this, we applied HyCo spheroids and invasion assays to testing the efficacy of P4HA1-inhibitor diethyl-pythiDC. By inhibiting collagen biosynthesis in tumor cells, diethyl-pythiDC directly limits their ability to remodel neighbouring microenvironment and thus to invade (53). As such, it has been proven to reduce tumor invasiveness preclinically, but has yet to reach clinical testing (53,64). Here, we confirmed the efficacy of diethyl-pythiDC on reducing invasiveness of STS117 HyCo spheroids, but could not do so for SK-LMS-1. Although advantageous, invasion assay must currently be performed off-chip due to Matrigel viscosity and limited on-chip well area. A chip-compatible invasion assay would require less viscous alternative matrix products or extensive chip design modification. Despite that, HyCo spheroids prove to be an adequate model for drug testing that could potentially improve the accuracy and success rate of bringing drugs from *in vitro* to *in vivo* to human trials.

Overall, we have demonstrated that our HyCo spheroids generated on-chip recapitulate clinical hallmarks of tumor hypoxia: gene signature of hypoxic STSs, hypoxia-induced treatment resistance and aggressive metastatic phenotype. Although described here for STSs, our device and methodology are inherently cancer agnostic and can be used with any spheroid-forming cell line of interest. Our ability to produce 15 HyCo spheroids per channel in only a few days, with high reproducibility, demonstrates that this technology can be efficiently used in a research setting. In addition, our microfluidic device is specifically designed to be as user-friendly and versatile as possible, broadening the scope of its possible uses and increasing its chances of adoption by the research community (65–68). At the fundamental level, our methodology can be applied to the investigation of response to hypoxia in cancer subtype-specific models, easily generating hypoxic gene expression data. As shown here, this data can in turn be used to find druggable targets, for which compounds can easily be tested on the very same clinically-relevant models using a variety of bioassays. By being complex enough yet easily-produced, HyCo spheroids demonstrate themselves to be a suitable *in vitro* model for researchers to investigate whether their findings can be converted into actionable targets, as well as model the downstream effects on phenotype before moving to *in vivo* studies. In addition, the radiocompatibility and broader user-friendliness of our microfluidic platform allows for easy investigation of treatment combinations, further increasing the potential of our technology and of our HyCo spheroids to bridge the gap between bench and benchmark in drug development.

## Materials and Methods

### Experimental design

The objectives of the study were as follows: on-chip generation of normoxic and HyCo spheroids from two human STSs cell lines, validation of hypoxia-specific gene expression in HyCo spheroids, validation of hypoxia-induced radioresistance and chemoresistance in HyCo spheroids, validation of increased invasiveness of HyCo spheroids, application of HyCo spheroids to drug testing based on RNAseq data. The following sub-sections describe the various methodologies associated to each of these objectives.

### Microfluidic chip

Negative molds of the top and bottom layers of the microfluidic chip were machined PMMA by computer numerical control Modela MDX-40A 3D milling machine (Roland, USA) using the chip design from our previously published paper. A mix of polydimethylsiloxane (PDMS) Dow SYLGARD 184 Silicon Elastomer Clear with a 1:10 ratio of Dow SYLGARD 184 curing agent (Ellsworth Adhesive, Canada) was used to cast the top and bottom layers of the chips. After a 15 min desiccation step, PDMS-filled molds were cured at 80 °C for 30 min in a Precision Compact oven (Thermo Fisher Scientific, Canada). The top and bottom layers of the chip were unmolded using tweezers, with inlets and outlets punctured using a 3 mm biopsy punch. Devices were then assembled manually after a 30 s exposure of each layer to atmospheric plasma using Enercon plasma gun (Enercon Industries Corporation, USA).

### Cell culture

SK-LMS-1 human leiomyosarcoma cell line was purchased from American Type Culture Collection (HTB-88, ATCC, USA). STS117 cell line was kindly provided by Dr. R. Gladdy (Mount Sinai Hospital, Canada). STS117 is a human STS primary cell line harboring a loss of function mutation of TP53, derived from patients’ primary extremity STS diagnosed as an undifferentiated pleomorphic sarcoma. SK-LMS-1 were cultured in Eagle’s minimum essential medium (EMEM, 320-005-CL, Wisent, Canada), supplemented with 10% fetal bovine serum (FBS, 12483-020, lot: 2567894RP, Gibco, ThermoFisher, Canada) and 1% penicillin-streptomycin mix (450-201-EL, Wisent, Canada). STS117 were cultured in Dulbecco’s modified Eagle medium/nutrient mixture F12 (DMEM/F12, 319-075-CL, Wisent, Canada), supplemented with 10% FBS and 1% penicillin–streptomycin solution. Both cell lines were maintained by subculturing at 80% of confluency for a maximum of 30 passages. Briefly, after the culture medium was aspirated, cells were washed with D-PBS and subsequently trypsinized with 0.25% trypsin EDTA (325-043-EL, Wisent Inc., Canada) for 3–5 min at 37 °C. Once detached, the enzymatic reaction was stopped by adding appropriately supplemented culture medium, and cells were passaged 1:4 in a new cell culture flask.

### Microfluidic Chip Preparation

Plastic inlets and outlets were connected to the microfluidic chips and assembled chips were placed in an empty pipet tips box for autoclave sterilization. Channels were first filled with 4 °C cold isopropanol to remove air bubbles. Beyond this step, the channels must remain filled with fluids to prevent air bubble formation. Then, the channels are washed three times with 200 µL 4 °C cold D-PBS (311-425-CL, Wisent, Canada), and three times with 4 °C cold PEG–PPG–PEG, Pluronic® F-108 solution (Sigma-Aldrich Canada Co, Canada). To prevent cell adhesion, Pluronic-filled devices were left to incubate overnight in 37 °C, 5% CO2 incubator. Prior to cell seeding, the channels are rinsed three times with 200 µL 37°C D-PBS and another three times with 200 µL of 37°C appropriately culture media.

### Spheroid Formation

To prepare cells for cell seeding, culture medium was aspirated, cells were washed with 5 mL of D-PBS and were trypsinized with 3 mL of 0.25 % trypsin 2.21 mM EDTA (325-043-EL, Wisent, Canada) and incubated for 3–5 min at 37 °C. Once detached, the enzymatic reaction was stopped by adding 7 mL of appropriately supplemented culture medium. The cell suspension was then collected in 50 mL Falcon tubes (Sarstedt Inc., Canada) and centrifuged for 5 min at 1500 rpm. The cell pellet is then resuspended in an appropriate volume to reach the final concentrations described in table 1. In prepared microfluidic devices, 200 µL of culture medium was aspirated from the outlet and 200 µL of cell suspension was added to the inlet. This step was performed three times and another three times starting from the outlet to the inlet, to ensure homogeneous distribution of the cell suspension. Culture medium is then changed every 24 hours until spheroids are formed.

**Table 1.** Cell suspension concentration for on-chip generation of HyCo and normoxic spheroids.

| Spheroid type | Concentration (cells/mL) |
| --- | --- |
| Hypoxic-Core (HyCo) spheroids | $4 \times 10^6$ |
| Normoxic spheroids | $2.5 \times 10^6$ |

### RNA extraction

Three days after seeding, 15 spheroids (1 channel) were washed on-chip by pipetting three times 200 µL of 37°C D-PBS (ref, Wisent, Canada), then, washed three times with 200 µL of RNAlater stabilization solution (AM7020, Invitrogen, ThermoFisher Scientific, Canada) directly on-chip. RNAlater stabilization solution was used to stabilize and protect cellular RNA of spheroids to avoid RNA degradation and to allow long-term storage of samples. Spheroids were retrieved from the chip in 800 µL of RNAlater, transferred in a 1.5 mL DNA LoBind Eppendorf tube (022431021, Eppendorf), 200 µL of PBS were added and tubes were kept overnight at 4°C. 600 µL of RNAlater-PBS buffer was removed, 400 µL of ice-cold PBS was added and sample were centrifuged at 4000 min^-1^.g for 3 min. The remaining 400 µL were removed, 600 µL of RLT buffer (Qiagen RNeasy® Plus Mini Kit, Cat. 74134, Qiagen, Canada) were added and tube was vortexed for 10 s. After, tubes were sonicated (xl-2000 Misonix, Canada) 3 times at level 6 for 1 s with 30 s rest on ice between each pulse. Lysates were centrifuge at 13000 min^-1^.g for 3 min. Supernatants were then transferred in gDNA eliminator spin columns placed in a 2 mL collection tubes. Next, samples were processed following Qiagen Purification of Total RNA from Animal Tissues protocol using Qiagen RNeasy® Plus Mini Kit. Of note, no β-mercaptoethanol was added to RLT buffer as RNAlater was already used to prevent RNA degradation. Finally, RNA concentration was measured using nano drop spectrophotometers (DS-11 FX, DeNovix, USA).

### RTPCR and qPCR

Reverse transcriptase (RT) PCR was performed using SuperScript IV VILO Master Mix with ezDNase enzyme kit (11766050, Invitrogen, Thermofisher, Canada) according to the manufacturer protocol. Mixture was gently homogenized, incubated at 37 °C for 2 min, briefly centrifuged and placed on ice. 4 µL of SuperScript IV VILO Master Mix and 6 µL of DNase-free water were added and gently mixed with the solution. Following steps were performed according to the manufacturer protocol. The resulting cDNAs were diluted 5-times in nuclease-free water and stored at -80°C until qPCR. PowerTrack SYBR green master mix kit (A46109, Applied Biosystems, ThermoFisher, Canada) was used for qPCR and samples were processed following the provided protocol. Samples were then placed in qPCR machine (Applied Biosystems™ StepOnePlus™ Real-Time PCR System). Expression of human GAPDH and Actin mRNA were used as endogenous control in the comparative cycle threshold method (2-*δδ*Ct) with the listed primers (PrimerBank, USA) (69–71).

### Bulk RNA Sequencing

Upon RNA extraction, samples were sent for sequencing at McGill Genome Center, Victor Phillip Dahdaleh Institute of Genomic Medicine at McGill. The following protocols were followed :

All samples had RNA integrity (RIN) measured with RNA TapeStation. All samples had RIN values between 6.9 and 9.4.

### RNA Sequencing Analysis

Unmapped paired-end sequences from an Illumina NovaSeq6000 sequencer were tested by FastQC v0.11.5 using a variety of metrics. Sequence adapters were removed and reads trimmed using Trimmomatic v0.39. The reads were mapped against the reference human genome (version hg38) using Burrows-Wheeler Aligner v0.7.17. Counts per gene were calculated with featureCounts v2.0.8 using annotation from Ensembl. Normalization and differential expression were calculated with DESeq2 (v1.46.0). Genes up- and down-regulated were identified with DESeq2 (p-value < 10e-6 and ≥ 2 fold change on pre-log2 transformed expression). Gene ontology terms enriched with protein-coding genes consistently induced by hypoxia in multiple cell lines were identified using WebGestalt 2024 (Benjamini corrected FDR < 0.05) (72).

### Radiotherapy treatment

Conventional EBRT was used for on-chip irradiation of HyCo and normoxic spheroids. Briefly, culture medium was replaced before whole-chip single dose irradiation using a Gammacell 3000 irradiator (Best Theratronics, Canada) at 4 or 8 Gy. Clonogenic assay was then performed immediately after irradiation following the previously described protocol.

### Drug treatment

#### Cisplatin

Cisplatin (1 mg/mL, DIN02355183, Accord Healthcare Inc., Canada) was obtained from the University of Montreal Hospital Center (CHUM) pharmacy and directly diluted in appropriate media at a final concentration of 5 µM (control: D-PBS). Treatment with cisplatin was performed 3 days after seeding by pipetting 3 x 200 µL in each channel. Then devices were incubated for 16 h at 37°C 5% CO2 incubator.

#### Treatment for invasion assays

Diethyl-pythiDC was obtained from MedChemExpress (HY-103068, USA) and resuspended in DMSO (MedChemExpress, USA) (Table 2). Treatment with diethyl-pythiDC was performed off-chip. Spheroids were retrieved and plated as previously described in the Invasion Assay section. Then, 90 µL of treatment-media solution was added on top of each spheroid and the plate was incubated overnight at 37°C 5% CO2 incubator. The next day, Invasion Assay protocol was performed and plate was put inside the Incucyte S3 for image acquisition.

**Table 2.**
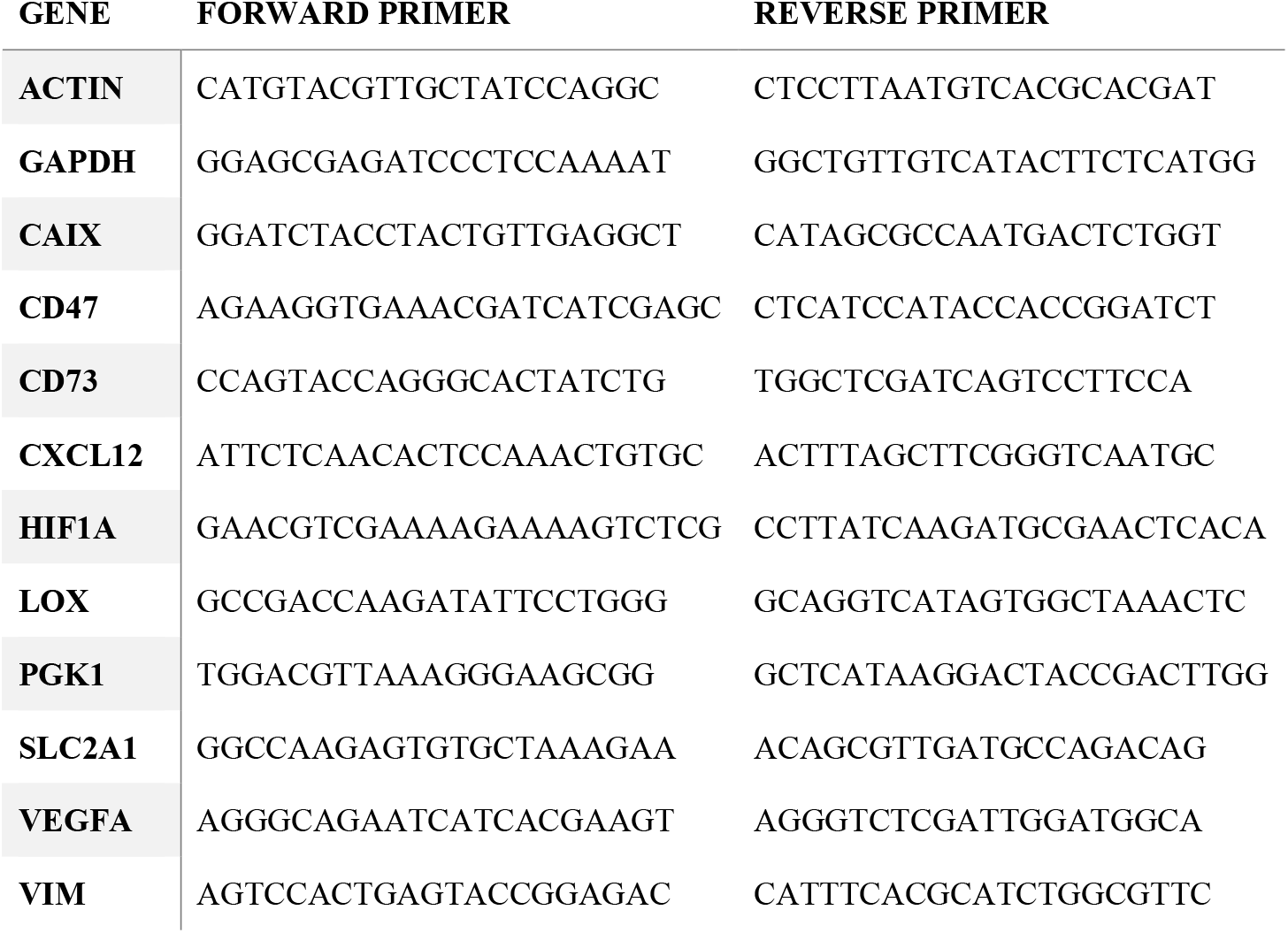
Primer sequences used for qPCR.

**Table 3.** Bulk RNA sequencing protocol.

|  |
| --- |
| QC, RNA, TapeStation |
| Library preparation RNAseq, NEBNextUltra II Directional RNA |
| Sequencing, Illumina NovaSeq6000 Sprime v1.5, PE100 (depth of ~400M reads per samples) |
| QC, Library for Illumina sequencing |

**Table 4.** Drug, buffer, concentration and control buffer concentration.

| Drug | Buffer | Concentration | Buffer Concentration (control) |
| --- | --- | --- | --- |
| Diethyl-pythiDC | DMSO | 50 μM | 1:2000 |

#### Immnunofluorescence Assay

For cisplatin treatment experiments, CellEvent Caspase-3/7 Green Ready Probe reagent was added to the culture medium of treated and untreated spheroids. 16 h after treatment, spheroids were washed three times with PBS directly in the device and fixed with Formalin 10% (Fisher Scientific Company, Toronto, ON, Canada) and 0.03 g/mL of sucrose (Invitrogen, USA) for 45 min. Then, spheroids were washed five times with PBS (Wisent Inc., Saint Jean-Baptiste, QC, Canada) and the top layer of the chip was peeled off. 15 spheroids per condition were retrieved using a cut-up 1000 µL pipet tip and pipetted directly into a histopathological cassette filled with optical cutting temperature (OCT) compound (Leica, Buffalo Grove, IL, USA). Included samples were left to sediment overnight, subsequently frozen on dry ice and stored at −80 °C. Frozen samples were sectioned using a Leica Cryostat (Leica, USA) with a 8 µm thickness and mounted onto glass slides. Sections were incubated for 1 h at room temperature with blocking buffer (PBS 1×, 3% IgG-free, Protease-free BSA, 0.5% Triton 100 10×). Sections were then incubated in a solution of blocking buffer with rabbit anti-CAIX (1:1000) (PA1-16592, ThermoFisher Scientific, USA) overnight at 4 °C. After washing three times with PBS, sections were then incubated in secondary antibody buffer (PBS 1×, 3% BSA) with AlexaFluor-647 antibody (1:750) (A31573, Invitrogen, USA) for 1 h at room temperature. Then, sections were stained with DAPI (1:5000 from 5 mg/mL stock solution) (D3571, Invitrogen, USA) to localize cell nuclei. Finally, sections were mounted using ProLong™ Gold Antifade Mountant (P36934, Invitrogen, ThermoFisher Scientific, USA). Fluorescence images were obtained on a Zeiss fluorescence microscope with ZEN microscopy software (Carl Zeiss AG, Germany).

### Clonogenic Assay

For each condition, after channels were washed with warm D-PBS, 5 HyCo spheroids and 21 normoxic spheroids were pipetted in a 15 mL Falcon tube. The spheroids were then centrifuged for 2 min at 1500 rpm, and D-PBS was aspirated. 300 µL of trypsin were added and left to incubate for 8 and 5 min respectively. Afterwards, a total of twenty ups and downs were performed with the pipette to ensure complete dissociation, and 700 µL of warm culture medium were added to block enzymatic reaction. 10 µL of cell suspension is then mixed to 10 µL of trypan blue in a 96-well plate, and 10 µL of this solution is pipetted in a cell counting slide to perform cell counting using a TC 20^TM^ Automated cell counter (BioRad, Canada). The cells are then centrifuged for 5 min at 1500 rpm, and the pellet is resuspended in appropriate volume to obtain a 1 × 10^5^ cells per mL solution. Then, the appropriate number of cells were seeded in triplicate in 6-well plates (3516, Corning Inc, Canada), of appropriate culture medium. Plates were kept in a 37°C 5% CO2 incubator with medium being changed every 3 days. After 10 days, wells were rinsed with 1 mL of D-PBS, and 2 mL 70% methanol solution containing 0.5% crystal violet (Sigma-Aldrich, Canada) was added to fix the colonies. After rinsing them with water, plates were imaged with Chemidoc (Bio-Rad, Canada). Colonies were manually counted using ImageJ software and normalized to the number of cells seeded to calculate clonogenic survival, which was in turn normalized to the control condition (73).

### Invasion Assay

Each microfluidic chip was placed in a petri dish to prevent spillage. The top layer was carefully peeled off using a pair of sterile tweezers, leaving the well layer containing the spheroids exposed. Then, 400 µL of appropriate culture media were added on top, and spheroids were harvested using a cut-up 1000 µL pipet tip to avoid damaging them. Spheroids are then individually pipetted in 90 µL of appropriate culture media into an ultra-low-attachment round bottom 96 well-plate (7007, Corning, USA). Media is then removed and 90 µL of media-drug mixture is added and spheroids are incubated at 37 °C 5% CO2 for 24h. The next day, the plate was cooled over ice using a cooling plate in a cool box until it reached approximately 4°C, at which point 90 µL of phenol-red free Matrigel (356237, Corning, USA) were added. The plate was then centrifuged at 300 rpm for 1 min at 4°C. Then, the plate was incubated for 30 min in a 37 °C 5% CO2 until the Matrigel polymerized. Finally, 50 µL of media alone or media-drug mixture were added, and the plate was placed in Incucyte S3 (Sartorius AG, Germany) for image acquisition over 10 days. Brightfield images were analyzed using custom ImageJ software analysis pipeline, by intensity thresholding to delimit three regions of interest (ROI) : edge, intermediate and core (Fig. 4.B) (73). Invasion indexes A and B were calculated using the following formulas. When comparing small normoxic and HyCo spheroids, invasion indexes A and B were normalized to their values at day 0 to account for size-induced biases. For each spheroid, invadopods were manually counted on the day where the difference in invasion indexes between conditions was the highest. Invadopod count is then normalized to the circumference of the edge ROI, estimated from the total invasion area, on the same day.

### Statistical analysis

Statistical analysis was conducted using GraphPad Prism (Version 9.0.1, GraphPad, USA). For spheroid size distribution, diameters were analyzed for significance using an unpaired t-test. For RNAseq, gene expressions were analyzed for significance using DESeq2 built-in statistical analysis. For gene ontology analysis, False Discovery Rate (FDR) correction is applied using Benjamini-Hochberg (BH) method. For clonogenic assay, clonogenic survival of spheroids were analyzed for significance using Holm-Šídák’s multiple unpaired t-test. For IF, caspase 3/7 signal was analyzed for significance using a two-way ANOVA with Šídák’s multiple comparisons test. For invasion assays, invasion indexes and invadopod densities were compared using a two-way ANOVA with Šídák’s multiple comparisons test and unpaired t-test, respectively. For invasion assay of treated spheroids, a mixed-effects analysis with Šidák’s multiple comparisons test was used due to uneven group sizes for SK-LMS-1.

## Acknowledgments

We thank Dr. Audrey Glory for her support and advice. We also thank Guillaume Cardin, Isabelle Clément, and Dr. Nicolas Malaquin for their help with EBRT irradiation. Finally, we thank Liliane Meunier and Véronique Barrès, of the CRCHUM’s molecular pathology core facility, for performing the cryo-sectioning and slide mounting steps.

## Author contributions

Conceptualization: ERM, PW, TG

Methodology: ERM, RC, JL, DY

Software: ERM, DY

Validation: ERM, RC, PW, TG

Formal Analysis: ERM, RC, DY

Investigation: ERM, RC

Resources: PW, TG

Data Curation: ERM, DY

Project administration: ERM, PW, TG

Visualization: ERM

Supervision: TG, PW

Funding acquisition: ERM, PW, TG

Writing—original draft: ERM, RC

Writing—review & editing: ERM, RC, JL, TG, PW

All authors have read and agreed to the published version of the manuscript.

## Data and materials availability

The data presented in this study are available on request from the corresponding author.

