## Supplementary Materials for "Hypoxic-Core (HyCo) Spheroids Recapitulate Hallmarks of Clinical Hypoxia: A Simple Chip-Based Method for Translational Oncology"

Elena Refet-Mollof *et al.*

**This PDF file includes:**

Figs. S1 to S4

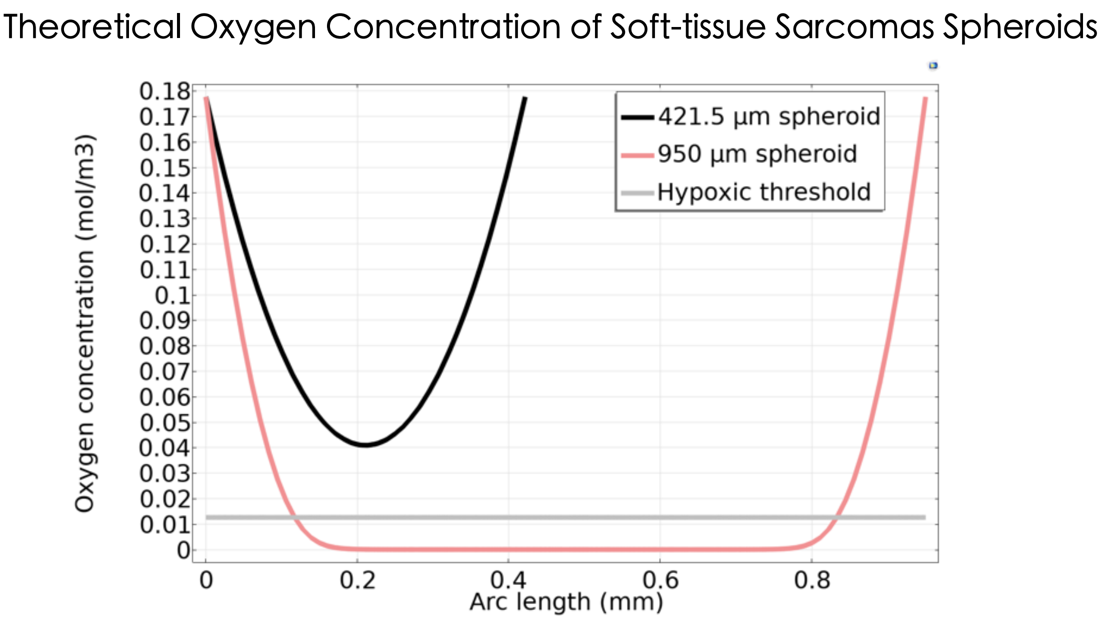


**Fig. S1.**

In silico model of the oxygen concentration in an on-chip soft-tissue sarcoma spheroid, depending on its diameter. HyCo spheroid diameter was set at 950 µm, and based on previous results normoxic spheroid diameter was set at 421.5 µm. Hypoxic threshold is set at 10 mmHg, i.e. the threshold for CAIX expression.

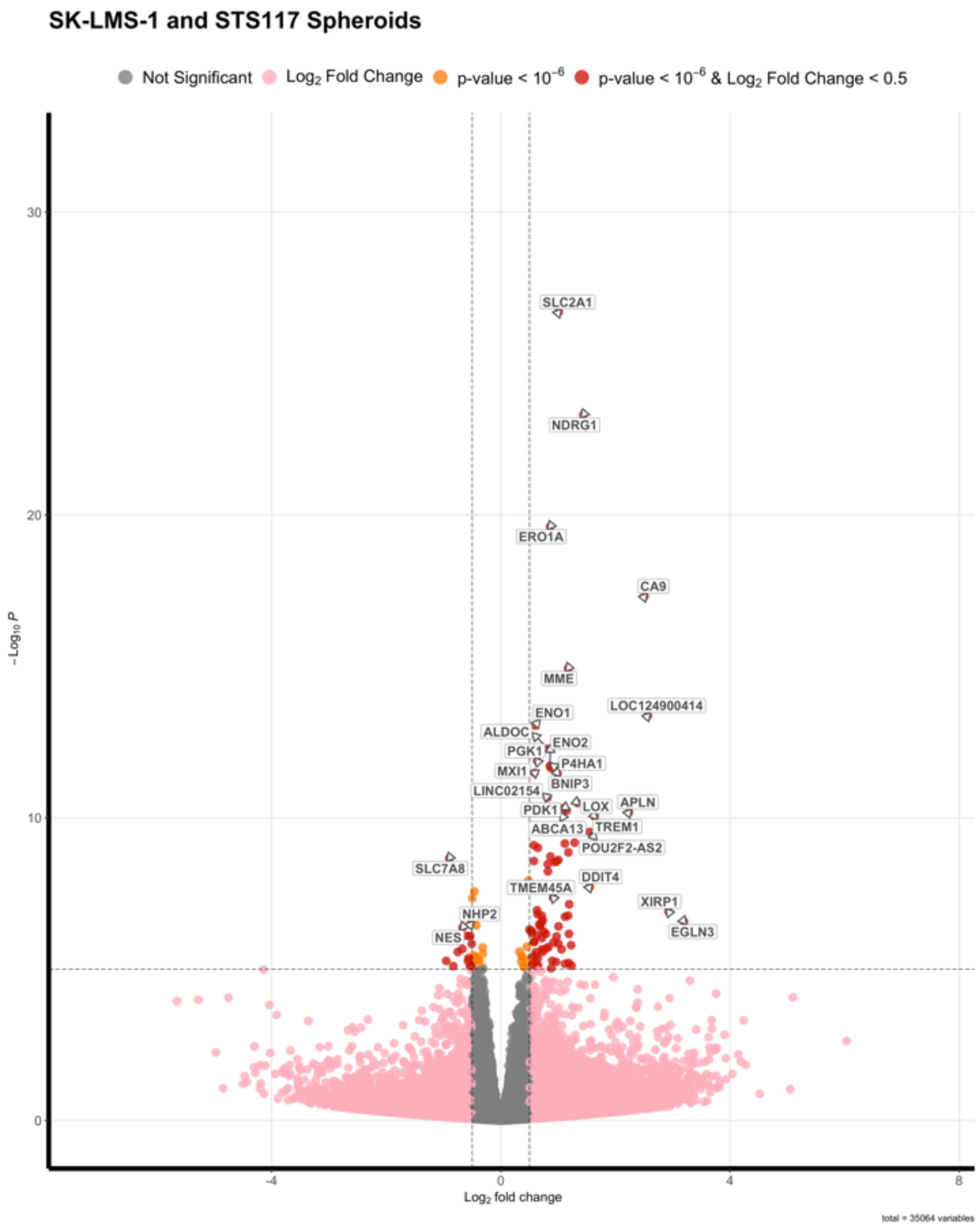


**Fig. S2.**

Volcano Plot of combined SK-LMS-1 and STS117 HyCo spheroids compared to normoxic spheroids. All data comes from N = 3 to 4 independent experiments.

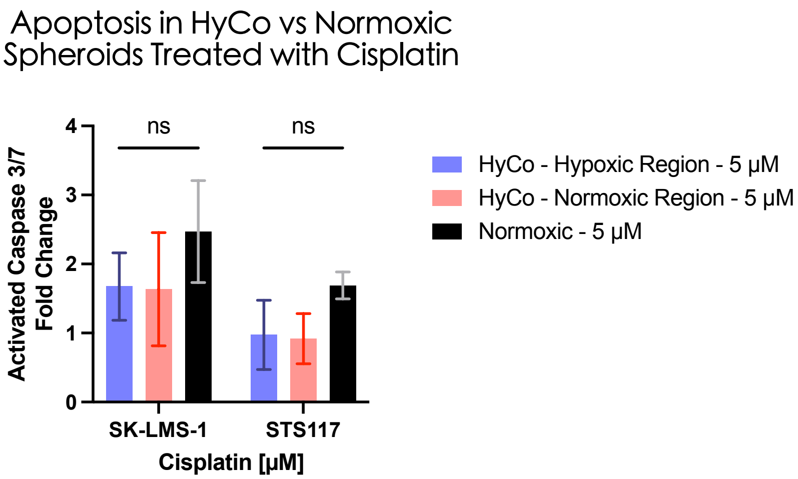


**Fig. S3.**

Quantification of apoptosis signal (activated caspase 3/7) after overnight exposure to 5μM of cisplatin in SK-LMS-1 and STS117 HyCo and normoxic spheroids. Data are presented as mean ± SD, (ns) : non-significant, N=3 independent experiments, n = 2 to 8 spheroids, 2way ANOVA, Šidák's multiple comparisons test.

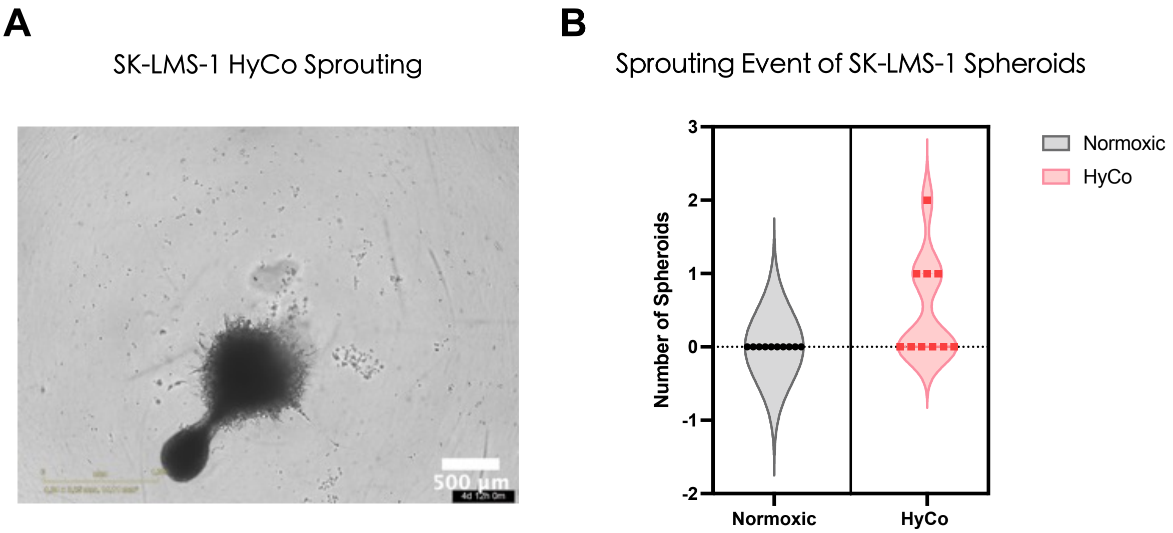


**Fig. S4.**

(A) Brightfield image of sprouting spheroid. (B) Quantification of sprouting event in SK-LMS-1 normoxic and HyCo spheroids over 10 days.
